# Clinical Antibiotic-Resistance Plasmids Have Small Effects on Biofilm Formation and Population Growth in *Escherichia coli* in vitro

**DOI:** 10.1101/2023.03.01.530568

**Authors:** Laura Brülisauer, Ricardo Leon-Sampedro, Alex R. Hall

## Abstract

Antimicrobial resistance (AR) mechanisms encoded on plasmids can affect other phenotypic traits in bacteria, including biofilm formation. These effects may be important contributors to the spread of AR and the evolutionary success of plasmids, but it is not yet clear how common such effects are for clinical plasmids/bacteria, and how they vary among different plasmids and host strains. Here, we used a combinatorial approach to test the effects of clinical AR plasmids on biofilm formation and population growth in clinical and laboratory *Escherichia coli* strains. In most of the 25 plasmid-bacterium combinations tested, we observed no significant change in biofilm formation upon plasmid introduction, contrary to the notion that plasmids frequently alter biofilm formation. In a few cases we detected altered biofilm formation, and these effects were specific to particular plasmid-bacterium combinations. By contrast, we found a relatively strong effect of a chromosomal streptomycin-resistance mutation (in *rpsL*) on biofilm formation. Further supporting weak and host-strain-dependent effects of clinical plasmids on bacterial phenotypes in the combinations we tested, we found growth costs associated with plasmid carriage (measured in the absence of antibiotics) were moderate and varied among bacterial strains. These findings suggest some key clinical resistance plasmids cause only mild phenotypic disruption to their host bacteria, which may contribute to the persistence of plasmids in the absence of antibiotics.

## 1. Introduction

Pathogens carrying antibiotic resistance (AR) plasmids are an increasing threat to public health, reducing treatment options for infected patients. But besides conferring resistance, AR plasmids can influence other phenotypic traits of their host bacteria, including biofilm formation (Burmølle et al. 2008; Dudley et al. 2006; Gallant et al. 2005; Gama et al. 2020; Ghigo 2001; May and Okabe 2008; Røder et al. 2013; Schaufler et al. 2016; Teodósio, Simões, and Mergulhão 2012; Yang, Ma, and Wood 2008). This matters in the context of infectious disease because biofilms are often involved in persistent infections (Costerton, Stewart, and Greenberg 1999; Lewis 2001; Stalder et al. 2020). Therefore, any phenotypic ‘side-effects’ of AR plasmids on biofilm formation may have downstream effects for bacterial epidemiological success and/or the likelihood of establishing persistent infections. More generally, plasmid-encoded changes in biofilm formation may be key determinants of evolutionary success, for both plasmids themselves and their host bacteria. In natural environments, bacteria very often live in biofilms, adhering to surfaces and embedded in an extracellular polymeric matrix (Flemming and Wuertz 2019; Hall-Stoodley, Costerton, and Stoodley 2004). This lifestyle can improve bacterial resistance to harsh environmental conditions (Lewis 2001), and biofilms may be hotspots for plasmid transfer via conjugation (Burmølle et al. 2008; Ghigo 2001; Røder et al. 2013; Virolle et al. 2020). Therefore, knowing whether and how plasmids affect bacterial biofilm formation would improve our fundamental understanding of the role of mobile genetic elements in bacterial evolution, and for clinically relevant AR plasmids this could increase our ability to predict and manage the spread of resistance in treatment contexts.

Despite both positive (Burmølle et al. 2008; Ghigo 2001; Reisner et al. 2006; Stalder et al. 2020) and negative (Gallant et al. 2005) examples, there is no consensus on how AR plasmids influence biofilm formation. One possible reason for this is that most studies to date only considered one plasmid and/or one bacterial host strain (Gama et al. 2020; Lim et al. 2010; Røder et al. 2013; Teodósio, Simões, and Mergulhão 2012). Given differences in strain construction, culture conditions, and other aspects of study design, this makes it hard to gain an overall view of how plasmids affect biofilm formation (i.e., to see the distribution of effect sizes). Therefore, individual studies testing multiple plasmids and strains simultaneously are required to estimate not only how plasmids affect biofilm formation, but how this varies among strains and plasmids. A second limitation of work to date is that many studies employed well-characterized laboratory plasmids and/or host strains (Burmølle et al. 2008; Gama et al. 2020; Ghigo 2001). Clinical plasmids and strains may behave differently compared to laboratory-adapted versions, for example due to carriage of genetic elements, such as prophages or other plasmids, that can interfere with acquisition of new plasmids and their downstream phenotypic effects (Gama et al. 2020; Igler et al. 2022). A third aspect, which also helps to explain the lack of consensus over plasmid effects on biofilm formation, is that there are several possible physiological mechanisms involved (Barrios Gonzalez et al. 2005; Gama et al. 2020). The variables playing a role include the AR genes and accessory genes encoded on the plasmid, but also the chromosomal genes of the bacterial host (Holden et al. 2021). Teasing apart the contributions of these different drivers therefore requires ‘swapping’ plasmids across bacterial strains, testing each plasmid in multiple strains and vice versa, which is often challenging with clinical strains/plasmids.

Here, we aimed to measure the effects of clinical AR plasmids on biofilm formation in *E. coli*, and to do this for several different bacteria-plasmid combinations. To achieve this, we used a combinatorial approach, swapping plasmids across host strains and quantifying their effects on biofilm formation. This allowed us to distinguish effects caused by differences among strains, plasmids and both together. For clinical relevance, we included five AR plasmids isolated from patients and belonging to the most problematic resistance plasmid families, ESBL, KPC and OXA-48 plasmids (Carattoli, 2009, Poirel et al., 2018). As bacterial host strains, we included clinical *E. coli* isolates obtained from hospitalized patients, the native hosts of some of our plasmids, and well-studied laboratory *E. coli* strains that differ genomically at known loci. We quantified plasmid-mediated effects on biofilm formation in two different sets of experimental conditions, motivated by past work indicating that biofilm formation in clinical *E. coli* isolates can depend on growth conditions (Naves et al. 2008). Finally, we also tested for a possible link between altered biofilm formation and plasmid effects on bacterial population growth in the absence of antibiotics. Our rationale for this part was that past work showed plasmids have variable effects on bacterial growth rate (Alonso-del Valle et al. 2021; Baltrus 2013; Rodríguez-Beltrán et al. 2021; San Millan and Maclean 2019; Vial and Hommais 2020), and the physiological drivers of these growth costs might also influence biofilm formation (Diaz Ricci and Hernández 2000). We found weak effects on biofilm formation for five clinical AR plasmids, specific to host background and nutrient condition. This provides new information about how AR plasmids influence biofilm formation and other phenotypic traits in epidemiologically successful, clinical bacteria.

## 2. Materials and Methods

### Plasmids

To test how natural, clinically relevant AR plasmids influence biofilm formation, we used five AR plasmids (pESBL1, pESBL15, pESBL25, pKPC-ecoli016 and a pOXA48-like plasmid; Table 1). These were obtained from *E. coli* isolates from hospitalized patients in two different studies at the University Hospital Basel, Switzerland (Noll et al. 2018; Tschudin-Sutter et al. 2016). For simplicity, we refer to pKPC-ecoli016 and pOXA-48-like as pKPC and pOXA-48 throughout the text. The plasmids pESBL1 (IncI, 111kb), pESBL15 (IncI, 88.9 kb) and pESBL25 (IncFIA, IncFIB, 131kb) encode extended spectrum *β*-lactamases (ESBL) of the CTX-M type. Plasmids pESBL1 and pESBL15 share a sequence similarity of 78% of coverage and 98.82% identity (Benz et al. 2020; Benz and Hall 2022; Tschudin-Sutter et al. 2016). The pESBL25 was not previously described and shows a mosaic structure sharing similarity with *E. coli* plasmids from the database (46%, 69% coverage and 99.77%, 99.81% identity with CP088793.1 and CP069982.1, respectively). The resistance genes encoded by pKPC and pOXA48 belong to the carbapenemases, a subgroup of *β*-lactamases. pKPC variants were previously found in *Klebsiella pneumoniae* (100% coverage and 99.42%, 99.86% with CP027700.1 and MF150120.1, respectively). The pOXA-48-like plasmid shares a 97% coverage and >99% identity both with the first described pOXA-48 plasmid (Poirel, Potron, and Nordmann 2012) and with pOXA48_K8, one of the best studied pOXA48-like variants (León-Sampedro et al. 2021).

**Table 1:** Resistance plasmids.

| Plasmid | Incompatibility group | Size (kb) | Resistance genes | Reference (Genbank Ac. No.) |
| --- | --- | --- | --- | --- |
| pESBL1 | IncI | 111 | *aadA5*, *bla*<sub>CTX-M-1</sub>,<br>*sul2*, *dfrA* | SAMN17073775 |
| pESBL15 | IncI | 88.9 | *bla*<sub>CTX-M-1</sub> | SAMN12275742 |
| pESBL25 | IncFIB, IncFIA | 131 | *bla*<sub>CTX-M-14</sub>, *tetB*,<br>*dfrA*, *mphA* | SAMN17073781 |
| pKPC | IncX3, IncU | 73.7 | *bla*<sub>KPC2</sub>, *sat2A* | UWWZ01000004.1 |
| pOXA48 | IncL | 63.6 | *bla*<sub>OXA48</sub> | UWXP01000003.1 |

### Bacteria and growth conditions

We used five *E. coli* strains (two clinical isolates and three laboratory strains) to investigate bacterial host-specificity of plasmid-mediated biofilm effects (Table 2). We included two clinical *E. coli* isolates, cESBL1 and cESBL15, cured of their native ESBL plasmids, originating from the clinical donor strains of the plasmids pESBL1 and pESBL15 (Benz et al. 2020). The clinical isolate cESBL1 belongs to the *E. coli* phylogroup G (sequencing type (ST) 117), which has been associated with poultry and various mammals as well as severe extra-intestinal diseases in humans and is characterized by high virulence and enhanced antibiotic resistance potential (Clermont et al. 2019; Lu et al. 2016). The second clinical isolate, cESBL15, is part of the B1 phylogroup in *E. coli* (ST40), a group adapted to a broad spectrum of hosts but predominantly found in vertebrate animals including humans and frequently harbouring extended-spectrum β-lactamases (Bajaj, Singh, and Virdi 2016; Berthe et al. 2013; Clermont et al. 2013; Martak et al. 2020). In addition, we used three isogenic laboratory strains, *E. coli* K-12 MG1655 (wild-type, hereafter referred to as MG1655), a variant thereof encoding a chloramphenicol resistance gene (*cat*) inserted into the *galK* gene locus (Δ*galK::cat*) and a chromosomal dTomato marker (hereafter referred to as MG1655+Cm), and another variant carrying a K43R mutation in the ribosomal *rpsL* gene, providing resistance to streptomycin (hereafter referred to as MG1655+Stm). We cultivated bacteria in lysogeny broth (LB) or in M9 minimal medium supplemented with 0.8% glucose and 1mM MgSO_4_. Overnight cultures prior to the experiments were supplemented with 100 mg/L of ampicillin to prevent plasmid loss, while no antibiotics were added during the biofilm formation assay.

**Table 2:**
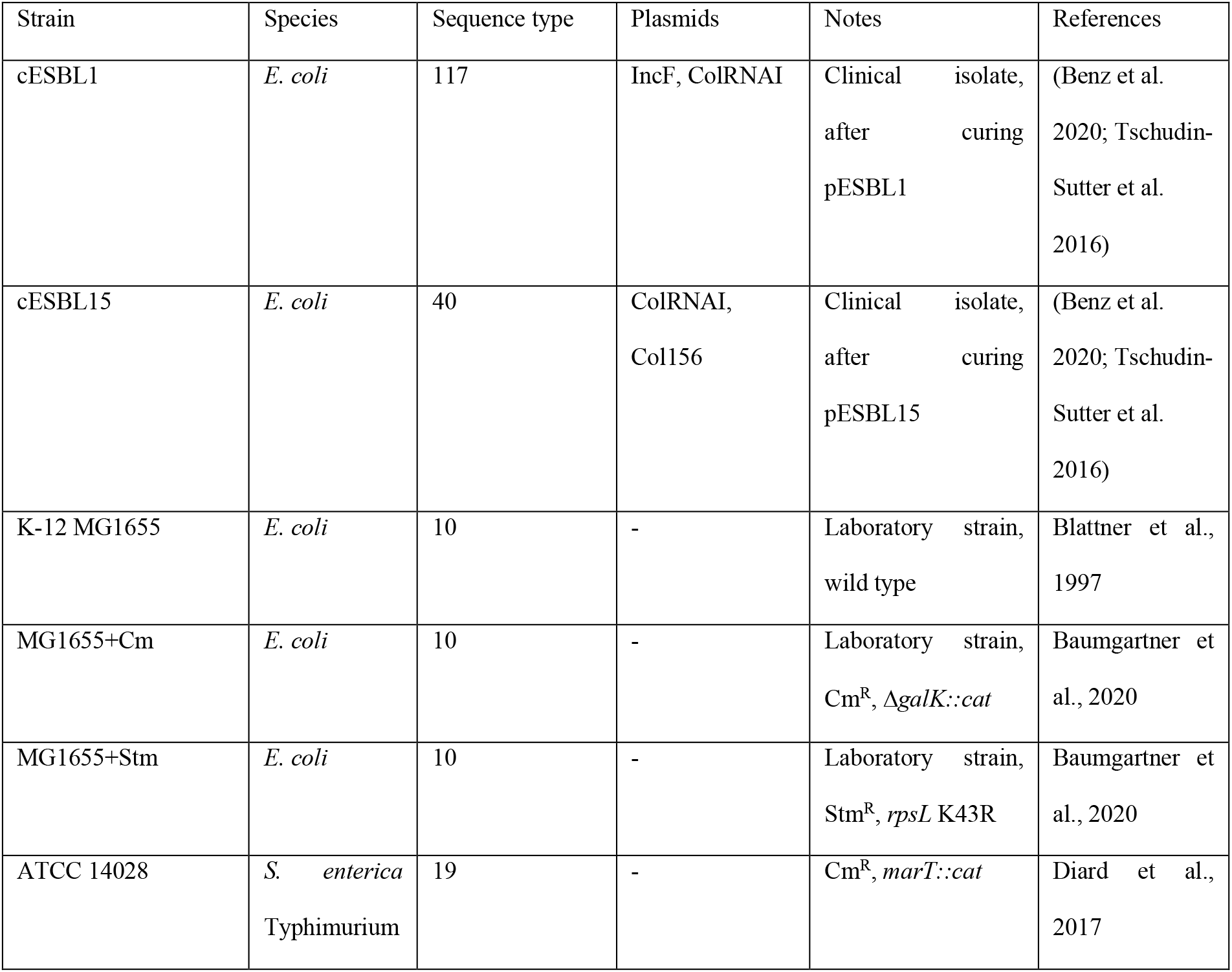
Bacterial host strains.

### Construction of novel plasmid-host combinations by conjugation

We introduced each of the five plasmids to each of the five bacterial host strains by conjugation (Alonso-del Valle et al. 2021), resulting in 25 different transconjugants. Briefly, we diluted overnight cultures (LB) of recipient and donor strains 1:100 and incubated them for 3.5h at 37°C. In late exponential growth phase, we harvested cells by centrifugation (1500 g; 15 min) and resuspended them in 0.9% NaCl, then mixed the recipient and donor cultures in a 1:1 (*v*:*v*) ratio. Subsequently, we plated serial dilutions on selective agar plates. For the two strains MG1655+Cm and MG1655+Stm, we obtained transconjugants by using selective plates containing ampicillin (100 mg/L) and chloramphenicol (25 mg/L) or streptomycin (100 mg/L), respectively. To obtain transconjugants of the wild-type MG1655 strain and the cured clinical donor strains, cESBL1 and cESBL15, we used a *Salmonella enterica* Typhimurium strain ATCC 14028 harbouring a chloramphenicol resistance marker (*marT::cat*) (Diard et al. 2017) as intermediate plasmid host, allowing us to discriminate *E. coli* transconjugants from *S. enterica* transconjugants on chromatic agar plates containing ampicillin (100 mg/L), because these three host strains do not carry any chromosomal resistance markers. We confirmed plasmid carriage and strain ID (for cESBL1 and cESBL15 transconjugants) by colony PCR. For colony PCRs, we first obtained template DNA by touching single colonies with a sterile pipette tip and suspending the cells in 20μL nuclease-free water. Then, we boiled the DNA at 95 °C for 15 min and added 2.5μL to 22.5μL PCR reagents. The PCR reaction mix contained 2x GoTaq® G2 HS Green master mix and 1 μM of the respective forward and reverse primer (see Table S1). Using a labcycler (Sensoquest, Göttingen, Germany), we ran the following thermal cycle program: initial denaturation for 4min at 95°C followed by 25 cycles of denaturation (30s at 95°C), annealing (45s at 65°C) and extension (1min at 72°C) and final extension at 72°C for 5min.

### Biofilm formation assays

We measured plasmid-mediated changes in biofilm formation for each plasmid-host combination in two sets of conditions: in LB medium and in M9 Minimal medium. We performed an adapted version of the crystal violet assay described by Christensen et al., 1985 and O’Toole et al., 1999. Briefly, we inoculated three single colonies of each strain in randomly assigned wells containing 200μL medium (LB or M9) of three separate 96-well microplates, resulting in 9 biological replicates of each of the 30 strains used (5 plasmids in each of the 5 hosts, plus 5 plasmid-free hosts). After overnight incubation at 37°C with agitation, using a 96-pin replicator, we inoculated test plates containing 180μL per well of the respective medium from the overnight cultures (1:180 dilution). After 24h static incubation at 37°C in a sealed plastic bag, we measured optical density at 600nm using a NanoQuant Infinite M200 Pro plate reader. Subsequently, we discarded the medium containing non-attached cells and washed the plates three times with 0.9% NaCl before staining with 0.1% crystal violet for 15min. Then, we repeated the three washing steps and air-dried the plates for 1h. We dissolved the retained crystal violet in 96% ethanol and measured absorbance at 595nm. We performed the biofilm assays in LB and M9 minimal media in separate experiments.

### Calculation of biofilm formation scores

We calculated three different measures of biofilm formation. First, “absolute biofilm formation” describes the amount of cells adhering to the polystyrene plate wells and is measured by absorbance after staining with crystal violet and re-dissolving in ethanol (OD595nm CV). Second, “normalized biofilm formation” gives the density of adhering cells relative to those in the liquid phase, calculated by dividing absolute biofilm formation (OD595nm CV) by the respective culture absorbance value (OD600nm culture). Third, to quantify the change in biofilm formation caused by the presence of the plasmid in a given strain-plasmid combination, we calculated “relative biofilm formation” using the following formula:

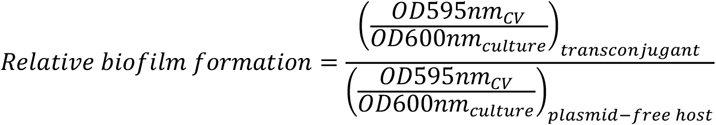

where values for the transconjugant come from a single microplate well (one replicate), and values for the plasmid-free host are the median for the corresponding plasmid-free strain (all replicates). We calculated the error associated with individual biofilm formation estimates using the following error propagation formula:

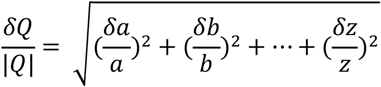

*a, b, …, z =* measurement values; *δ* = error associated with measurement value; Q = calculated value from measurements a, b, …, z (in this study for example relative biofilm formation)

### Growth curves and plasmid effects on bacterial growth

We determined plasmid growth effects in all hosts in LB medium and M9 Minimal medium by monitoring growth in liquid medium in the absence of antibiotics in microplates over 24h, then comparing transconjugants with their corresponding plasmid-free hosts. From a subset of the overnight cultures used for the biofilm formation assay above, we diluted three biological replicates of each strain into fresh medium using a pin replicator and incubated for 24h at 37°C in a plate reader (Tecan NanoQuant Infinite M200 Pro). We measured absorbance at 600nm every 15min after 5s of shaking (amplitude = 1.5mm). We determined the maximum growth rate (μmax), maximum optical density (OD_max_ and area under the curve (AUC) using the growthrates (Thomas Petzoldt 2020) and flux (Jurasinski et al. 2022) packages in R, version 0.8.2. Further, we calculated the relative maximum growth rate by dividing the maximum growth rate of each transconjugant by the maximum growth rate of the respective plasmid-free host (as for relative biofilm formation). In this case, due to normally distributed data, we used the arithmetic mean. We calculated the errors using error propagation as above.

### Statistical analysis

We tested for variation of biofilm formation (relative, normalised or absolute; see above) among strains and plasmids using a two-way ANOVA. We tested the significance of individual effects (the difference between plasmid-carrying and plasmid-free versions in each strain-plasmid combination) by Welch’s two-sample *t*-test using the Holm-Bonferroni correction for multiple testing (Dunn 1961; Holm 1979; Neyman and Pearson 1928). To estimate effect sizes as percentage changes in biofilm formation relative to plasmid-free strains, we took the average relative biofilm formation score in each combination, converted it to a percentage change (subtracting 1 then multiplying by 100), before removing negative signs (by squaring and rooting each value) to give all values as a percentage deviation in either direction.

## 3. Results

### Weak effects of clinical antibiotic resistance plasmids on biofilm formation

Across the 25 combinations of five clinical plasmids (belonging to four different incompatibility groups) and five bacterial strains we tested in nutrient-rich LB medium (Fig. S1), changes in biofilm formation caused by plasmid introduction were on average weak (Fig. 1A; mean relative biofilm formation=1.14, s.d.=0.46). This amounted to an average effect size of ±18.61%, s.d.=39.31 (in either direction, calculated from the squared and rooted mean percentage change in each combination). Despite this weak average effect across the dataset, there were some relatively strong individual effects (Fig. 1B), with a magnitude that varied depending on both the plasmid (two-way ANOVA: F_4,200_=40.67, p<0.001), the host strain (F_4,200_=19.15, p<0.001), and their interaction (F_16,200_ = 29.76, p < 0.001). Specifically, there were only three plasmid-host combinations where we detected significant changes in biofilm formation upon plasmid introduction (tested by Welch’s two-sample *t*-test with Holm-Bonferroni correction, Fig. 1B). The strongest such effect was an increase by 214% in biofilm formation induced by plasmid pESBL25 in the clinical isolate cESBL15 (Fig. 1B). This combination was strikingly different from our observations in other combinations, so we carried out an additional experiment using six independently constructed clones of this plasmid-host combination. This revealed no such increase in biofilm formation (Fig. S2), suggesting the outlying measurement in the main dataset was specific to that particular assay and, therefore, this individual observation should be interpreted with caution. If we exclude this combination from the above analyses, the qualitative outcome is unchanged (effects of host strain, plasmid and interaction remain significant at p<0.001), although the average effect size drops to ±11.06%, s.d.=11.17. The other two significant changes we detected upon plasmid introduction were both in strain MG1655+Stm, with plasmids pKPC and pOXA48 (Fig. 1). In summary, plasmids did not significantly alter biofilm formation in most combinations, and in the few cases where they did, these effects were specific to particular plasmid-host combinations.

**Figure 1:**
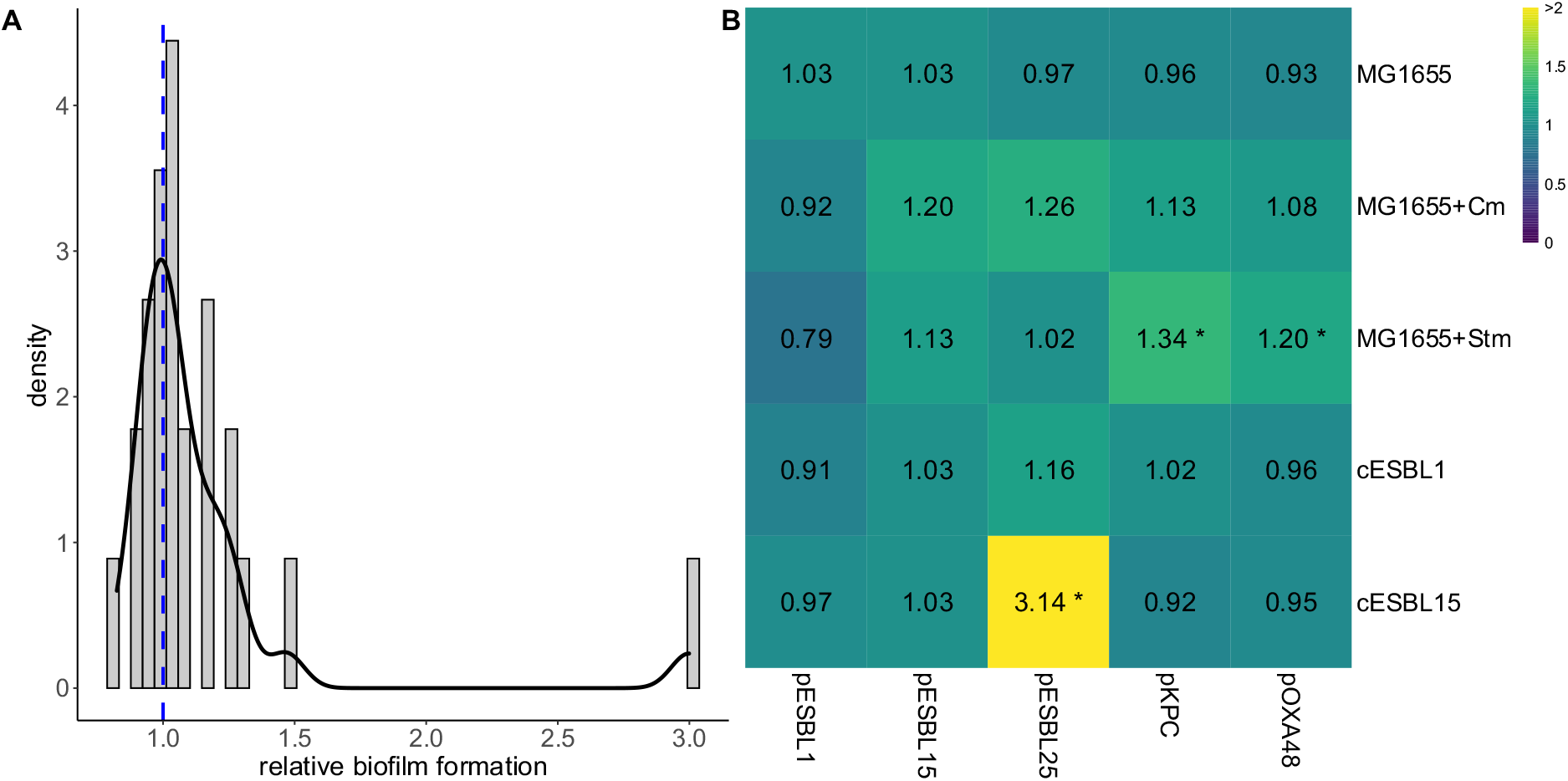
A: Distribution of relative biofilm-formation scores (plasmid-carrying vs plasmid-free) for 25 different bacterium-plasmid combinations in LB medium. The score for each combination is taken as the median from nine replicate assays (see Methods); a score of 1.0 indicates no change in biofilm formation upon plasmid introduction. B: Variation of relative biofilm formation scores among the five host strains (columns) and five plasmids (rows); as in the left-hand panel, each score is the median from nine replicates (normalised and absolute scores for all replicates are provided in Fig. S1 and S4). Asterisks indicate combinations with p < 0.05 (Welch’s two-sample t-test comparing plasmid-carrying vs plasmid-free, corrected for multiple testing using the Holm-Bonferroni method). Note the colour scale does not cover the whole range of data.

We found further evidence that host genetic background matters for biofilm formation by comparing the five plasmid-free host strains: their normalised biofilm formation scores of the plasmid-free strains varied significantly (one-way ANOVA: F_4,40_= 55.40, p<0.001), with MG1655+Stm scoring highest. This shows the K43R mutation in the ribosomal *rpsL* gene (carried only by MG1655+Stm) promoted biofilm formation in plasmid-free *E. coli*. Furthermore, plasmid-carrying strains in this bacterial host consistently had higher absolute and normalised biofilm scores (Fig. S1, S3, S4), showing the effect of the *rpsL* mutation on biofilm formation was maintained after plasmid acquisition.

### Similar, weak effects of plasmids on biofilm formation in minimal growth medium

We tested whether our above results were specific to LB medium by carrying out a separate biofilm assay in M9 minimal medium. This revealed a similar pattern to that observed above in LB medium (Fig. 2): on average, plasmid introduction did not lead to large differences in biofilm formation compared with corresponding plasmid free strains (Fig. 2A; mean relative biofilm formation = 1.29±1.38, mean effect size =37.67%, s.d=135.74). We detected a significant individual effect only for the same anomalous combination as above (cESBL15 with pESBL25; Fig. 2B). Repeating the analysis without this combination, the average effect size decreased to ±10.57%, s.d.=8.01, and there was no significant variation of average effects among plasmids (two-way ANOVA: F_4,192_=0.40, p>0.05), although as above in LB there was variation among host strains (two-way ANOVA: F_4,192_=9.61, p<0.001) and the strain × plasmid interaction (two-way ANOVA: F_15,192_=2.16, p<0.01). As above in LB, strain MG1655+Stm was the strongest biofilm former among plasmid-free strains (one-way ANOVA on normalised biofilm formation scores: F_4,40_=129.50, p<0.001), and remained so after plasmid acquisition. In summary, the experiment in minimal medium was consistent with our findings in rich medium, showing that plasmids had only moderate effects on biofilm formation in most combinations, and supporting the effect of the *rpsL* mutation carried by MG1655+Stm on biofilm formation.

**Figure 2:**
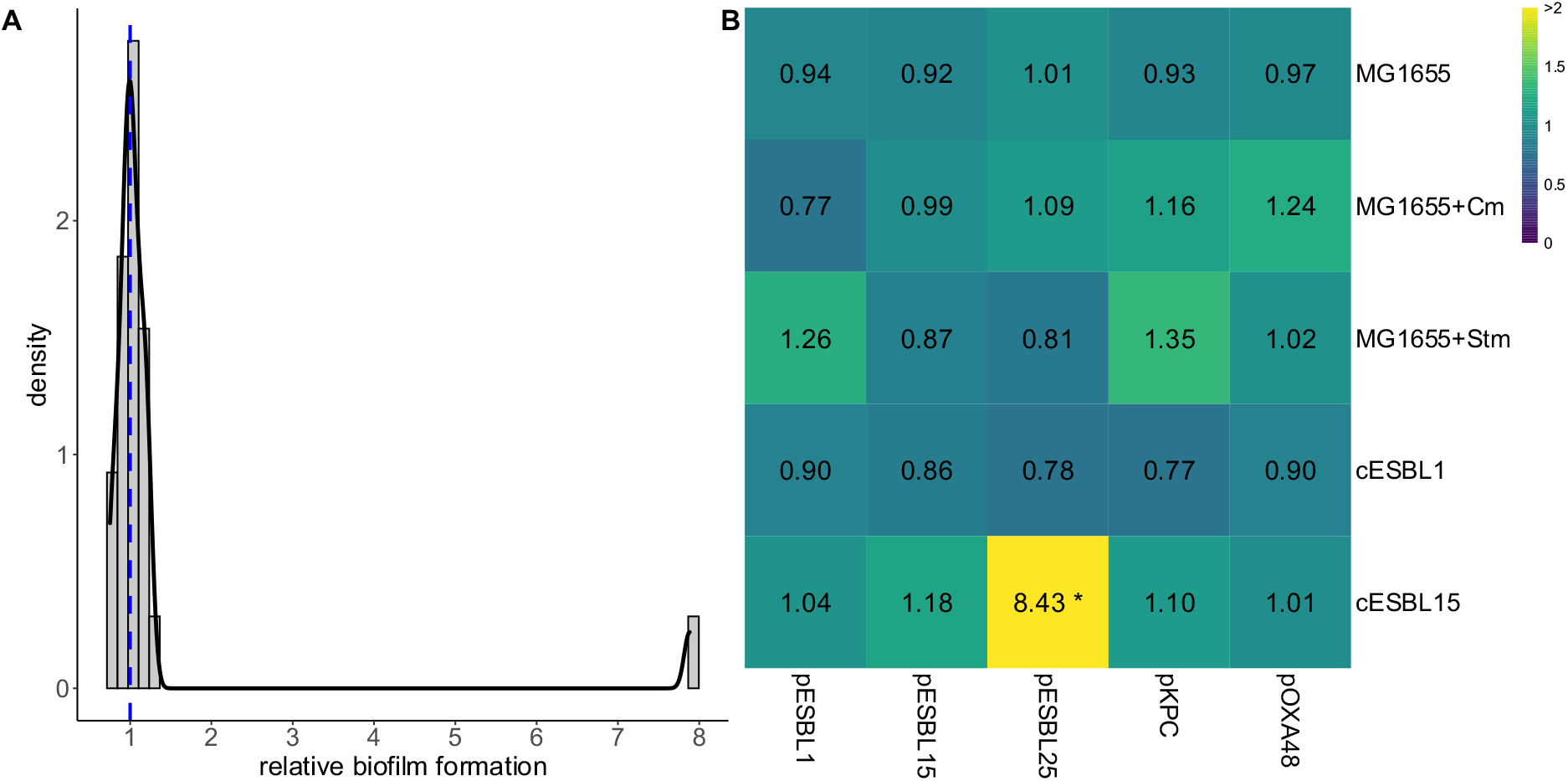
A: Distribution of relative biofilm-formation scores (plasmid-carrying vs plasmid-free) for 25 different bacterium-plasmid combinations in M9 medium. The score for each combination is taken as the median from nine replicate assays (see Methods); a score of 1.0 indicates no change in bioßfilm formation upon plasmid introduction. B: Variation of relative biofilm formation scores among the five host strains (columns) and five plasmids (rows); as in the left-hand panel, each score is the median from nine replicates (normalised and absolute scores for all replicates are provided in Fig. S3 and S4). Asterisks indicate combinations with p<0.05 (Welch’s two-sample t-test comparing plasmid-carrying vs plasmid-free, corrected for multiple testing using the Holm-Bonferroni method). Note the colour scale does not cover the whole range of data.

### Moderate, host-dependent effects of plasmids on bacterial growth rate

We measured population growth rates in the absence of antibiotics for all 25 plasmid-host combinations and their plasmid-free equivalents. In most cases, we detected no significant effects of plasmid acquisition on growth rate (Fig. 3). The average relative maximum growth rate in LB medium was 0.94, s.d.=0.08, and in M9 medium 0.99, s.d.=0.17 (Fig. 3). The five host strains paid different average growth costs upon plasmid acquisition (effect of strain in two-way ANOVA: F_4,50_=3.08, p<0.05), although the strength of this effect depended on the plasmid (strain × plasmid interaction: F_16,50_=2.55, p<0.01). For example, strain MG1655 paid relatively large costs for some plasmids such as pESBL15, but not for pESBL25 (Fig. 3A). Despite this, different plasmids were not associated with different growth costs on average (F_4,50_=1.34, p>0.05). In M9 minimal medium there was also variation among host strains (F_4,50_=6.78, p<0.001), plasmids (F_4,50_=2.58, p<0.05), but no strain × plasmid interaction (F_16,50_=1.44, p>0.05). The plasmid-bacterium combination cESBL15 with pESBL25 paid the largest growth cost in LB and in M9 (Fig. 3B). This was the same combination where we observed anomalous biofilm formation scores above. From the same dataset, we can extract other growth parameters including the maximum population density (maximum OD) and the area under the curve (AUC). These parameters provide a similar picture compared to growth rate, with moderate effects that varied among host strains (Fig. S5, S6). Finally, we tested for an overall association between growth costs and biofilm formation across the 25 strain-plasmid combinations, but found no evidence of this (Fig. S7). In summary, changes to bacterial growth rate in the absence of antibiotics upon plasmid acquisition were mostly weak, but depended on the bacterial host strain.

**Figure 3:**
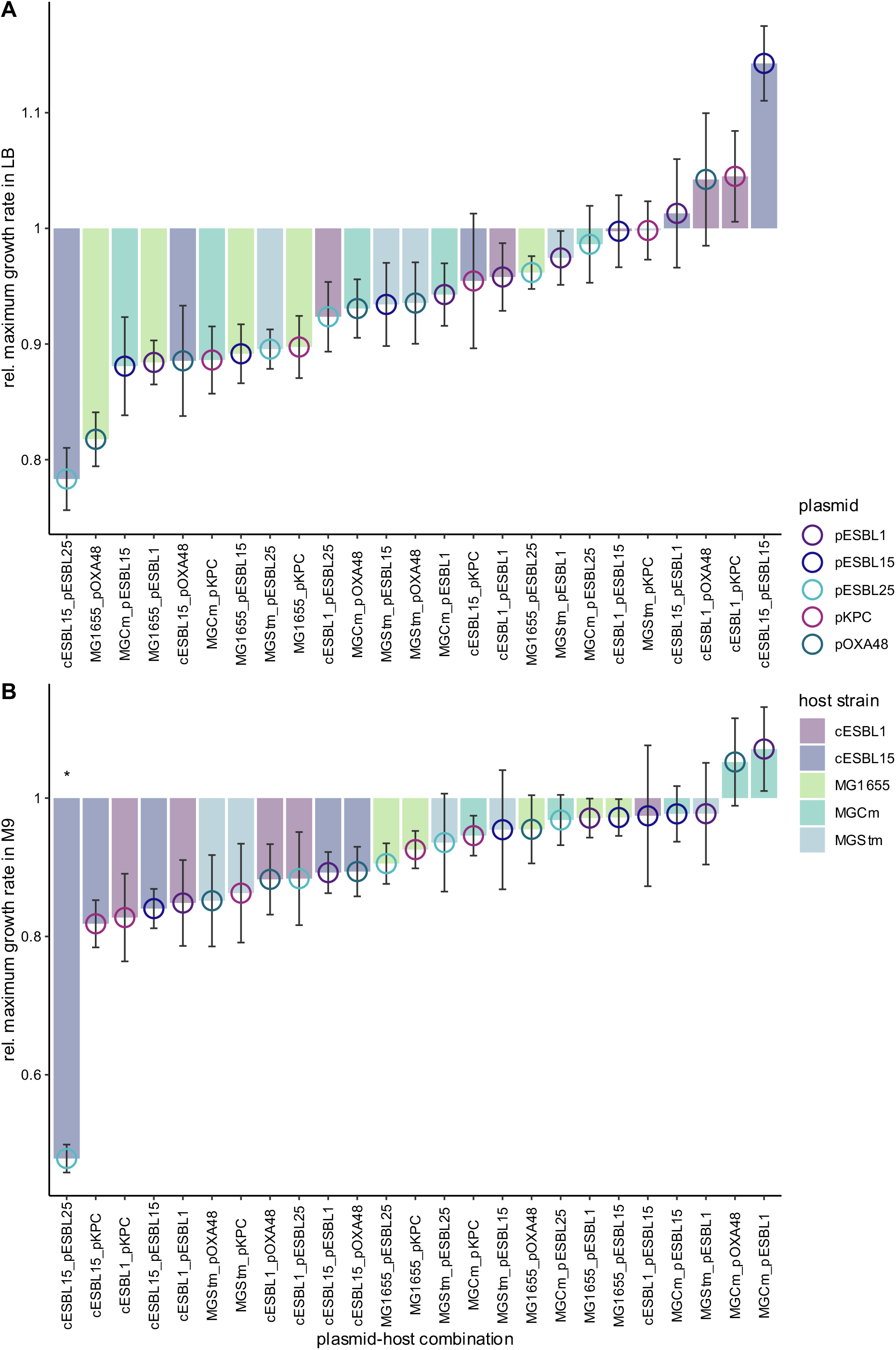
Relative growth rates (of plasmid-carrying strains relative to plasmid-free equivalents) in all combinations in LB medium and M9 minimal medium. (n=3). Cases with p<0.05 (Welch’s two sample t-test comparing plasmid-carrying vs plasmid-free, corrected for multiple testing using the Holm-Bonferroni method) are indicated with an asterisk. Error bars indicate propagated standard error.

## 4. Discussion

In a combinatorial screen, we studied the effects of clinical AR plasmids on biofilm formation and growth rate in various bacterial hosts and in two nutrient conditions. Our experimental design enabled us to separate the contribution of the plasmid and the host strain to biofilm and growth effects. This revealed plasmid effects to be specific to particular plasmid-host combinations. However, the overall distribution of plasmid effects across our dataset showed such effects to be in general moderate: clinical plasmids did not significantly alter biofilm formation in most combinations, and when they did their effects were weak. This contrasts with several past observations of strong plasmid-mediated changes in biofilm formation in studies with smaller numbers of plasmid-bacterium combinations or laboratory strains (Burmølle et al. 2008; Dudley et al. 2006; Gallant et al. 2005; Gama et al. 2020; Ghigo 2001; Røder et al. 2013; Stalder et al. 2020; Teodósio, Simões, and Mergulhão 2012; Yang, Ma, and Wood 2008). Consistent with the moderate phenotypic effects in our biofilm dataset, plasmids also had only weak effects on bacterial growth in the absence of antibiotics, agreeing with the findings of Alonso-del Valle et al. (2021), who reported weak growth effects of a clinical pOXA48 plasmid in natural enterobacterial strains. Moreover, our finding of a weak average growth cost here aligns very well with a meta-analysis by Vogwill and Maclean (2015). In their analysis of 77 publications, Vogwill and Maclean (2015) found an average relative growth cost of 0.91±0.024 for AR plasmids, across studies employing different methodologies and fitness measures. The average relative growth rate in our LB dataset of 0.94±0.08 is comparable to this. Thus, our combinatorial approach revealed host-dependent, but on average small changes in growth rate and biofilm formation upon plasmid acquisition.

The key strength of this study is that we used a fully factorial design with clinical AR plasmids and a combination of clinical and laboratory host strains, going beyond past work studying laboratory plasmids and / or host strains. Clinical host strains differ from model strains because they can carry additional genetic elements such as plasmids, prophages and pathogenicity islands, which can potentially interfere with the phenotypic effects of new plasmids (Gama et al. 2020; Igler et al. 2022). Nevertheless, having included three laboratory strains as well, we found a similar picture in both clinical and laboratory strains. This suggests the moderate effects we observed for these clinical plasmids are likely to apply in other strains as well. Furthermore, our experimental design allowed us to disentangle the contributions of plasmid and host strain to the biofilm effect. This showed both factors shape biofilm effects, with the host strain having a relatively strong influence. A key implication of this is that plasmid effects on bacterial ecology can be specific to the plasmid-host combination. Thus, predicting the spread of plasmid-encoded resistance genes and their downstream effects on biofilm formation requires information not only about which plasmids are circulating, but which strains they are carried by.

Our results contrast with several past observations of plasmids affecting biofilm formation (Burmølle et al. 2008; Gallant et al. 2005; Ghigo 2001; Reisner et al. 2006; Schaufler et al. 2016; Yang, Ma, and Wood 2008). One possible reason for this, which could potentially be explored in future work, is the role of mating-pair formation and conjugation itself. Some past studies showed strong effects of plasmids on biofilm formation upon co-inoculation with viable plasmid recipient strains (Ghigo 2001; Reisner et al. 2006; May and Okabe 2008). Ghigo (2001) proposed a key role for pilus expression, followed by conjugation and transient derepression of pilus synthesis in new transconjugant cells. In our experimental set-up, with pure cultures of plasmid-carrying strains, there are no/fewer recipient cells than in such co-inoculation experiments, and transfer to other plasmid-carrying cells is probably limited by surface exclusion (Achtman and Kennedy 2023). Consistent with this, we identified putative entry exclusion system genes for all the plasmids tested here (Table S2). The biofilm effects we observed are therefore more likely to stem from the plasmid’s interaction with the bacterial host’s gene expression and metabolism (Barrios Gonzalez et al. 2005; Gama et al. 2020). Nevertheless, we do not rule out that these plasmids would have stronger effects in other conditions where conjugative transfer is more common, particularly because these plasmids are conjugative (Benz et al. 2020; Pathak et al. 2022; Teodósio, Simões, and Mergulhão 2012). Crucially, Ghigo (2001) also observed plasmid effects on biofilm formation for several plasmids in pure cultures, and other past studies have also used a pure-culture set-up as we did (Burmølle et al. 2008; Gallant et al. 2005; Schaufler et al. 2016; Yang, Ma, and Wood 2008). Therefore, the lack of strong effects in our experiments are not fully explained by the pure culture set-up.

In contrast to the mild effects we observed for plasmid-borne resistance, we found a strong effect of a chromosomal resistance mutation (*rpsL* K43R in *E. coli* K-12 MG1655), resulting in a significantly higher baseline biofilm formation which was maintained after plasmid acquisition. This effect is consistent with previous studies linking *rpsL* mutations to pleiotropic changes in other bacterial phenotypes (Paulander, Maisnier-Patin, and Andersson 2009) and biofilm formation in particular (Boehm et al. 2009). More generally, chromosomal resistance mechanisms frequently affect essential and/or conserved genes, such as ribosomal genes like *rpsL* and those encoding RNA polymerase subunits, often leading to wide-ranging effects on expression of other genes and phenotypes (Hall et al. 2015; Perkins and Nicholson 2008; Qi, Preston, and Maclean 2014). The relatively strong side effects we observed for a chromosomal resistance mutation compared to plasmid-borne resistance may therefore be consistent with a wider trend that plasmid-borne resistance comes with relatively few pleiotropic effects compared to, for example, resistance mutations affecting ribosomal subunits or RNA polymerase. We speculate further that relatively mild effects of clinical resistance plasmids on both biofilm formation and population growth may contribute to their stability in the absence of antibiotics, in that bacteria can acquire them without incurring large costs. This is also consistent with past work finding a similarly low cost of plasmid-borne resistance (Vogwill and Maclean (2015); see also Alonso-del Valle et al. (2021)). Note this does not exclude the idea that plasmids which increase biofilm formation may gain an evolutionary advantage, due for example to increased opportunities for horizontal transfer and improved survival of their bacterial hosts (Burmølle et al. 2008; Ghigo 2001; Røder et al. 2013). Some plasmids may be under positive selection for such effects in some conditions, but our experiments suggest strong changes in biofilm formation are not general, in that they did not arise in most of the plasmid-bacterium combinations we tested in vitro.

We were able to visualise the distribution of plasmid effects on biofilm formation across 25 combinations in two different experimental conditions, going beyond past work with smaller numbers of strains and in non-combinatorial designs. Our dataset nevertheless has some limitations. First, the number of significant effects in our dataset was small. In a larger screen with even more combinations, a clearer picture could be obtained of the variance and mean effect among the subset of combinations where plasmids do alter biofilm formation. We note there is a significant practical hurdle to scaling up the combinatorial approach used here, in terms of plasmid curing and re-introduction effort increasing as more strains are added. Second, one strong biofilm effect (combination cESBL15::pESBL25) was not reproducible in further validation experiments (Fig. S2). This emphasizes that individual observations should be interpreted cautiously. Despite this, our key findings (moderate effects on average, specificity for particular combinations) emerge from the entire dataset across the different combinations, rather than from individual datapoints. Also, the results were similar across both nutrient conditions. Therefore, our key findings are likely robust.

In conclusion, we find weak effects of clinical resistance plasmids on biofilm formation and population growth in a range of clinical and laboratory strains. This suggests clinical AR plasmids affect bacterial phenotypes less than might be expected from previous studies with different study designs, but in a way consistent with recent studies detecting moderate effects of resistance plasmids on bacterial population growth (Alonso-del Valle et al. 2021; Vogwill and Maclean 2015). This in turn implies that plasmids may incur only mild costs in terms of disruption of other phenotypes, which potentially contributes to their persistence in the absence of antibiotic selection.

## 5. Acknowledgements

We thank Fabienne Benz for advice and for providing plasmid-free versions of clinical strains, Adrian Egli for providing the original clinical strains, and Gregory Velicer for feedback. Funding: Swiss National Science Foundation project number 310030_192428.

## 6. Competing interests statement

We have no competing interests to declare.

## Supplementary Material

### Supplementary Figures

**Supplementary Figure 1:**
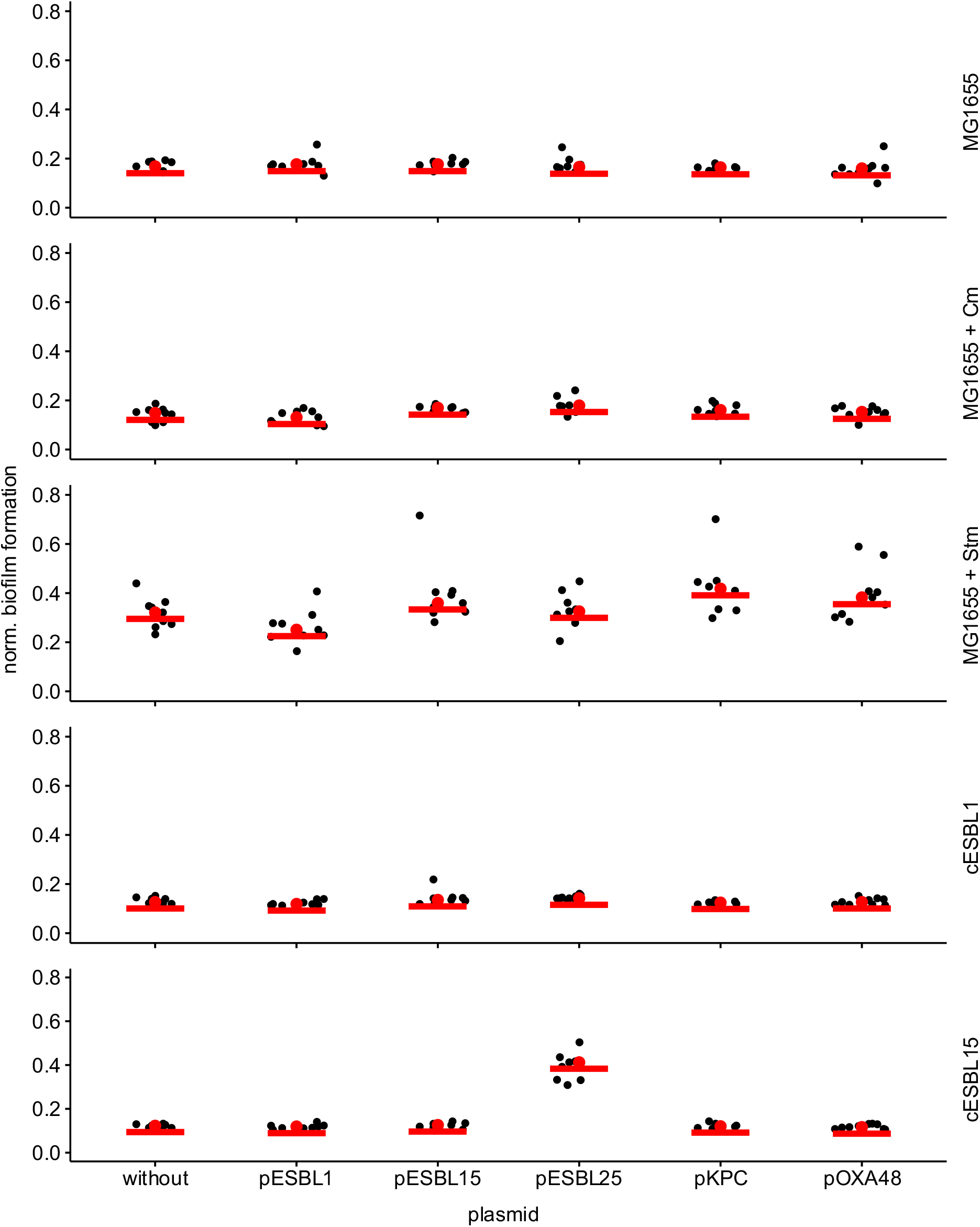
Normalised biofilm formation in LB medium for 25 plasmid-bacterium combinations, including plasmid-free reference strains. Each panel shows the biofilm formation in one host strain (given by the label at right) with each of the five plasmids or the plasmid-free version (x-axis). Points show normalised biofilm formation in each replicate (n = 9).

**Supplementary Figure 2:**
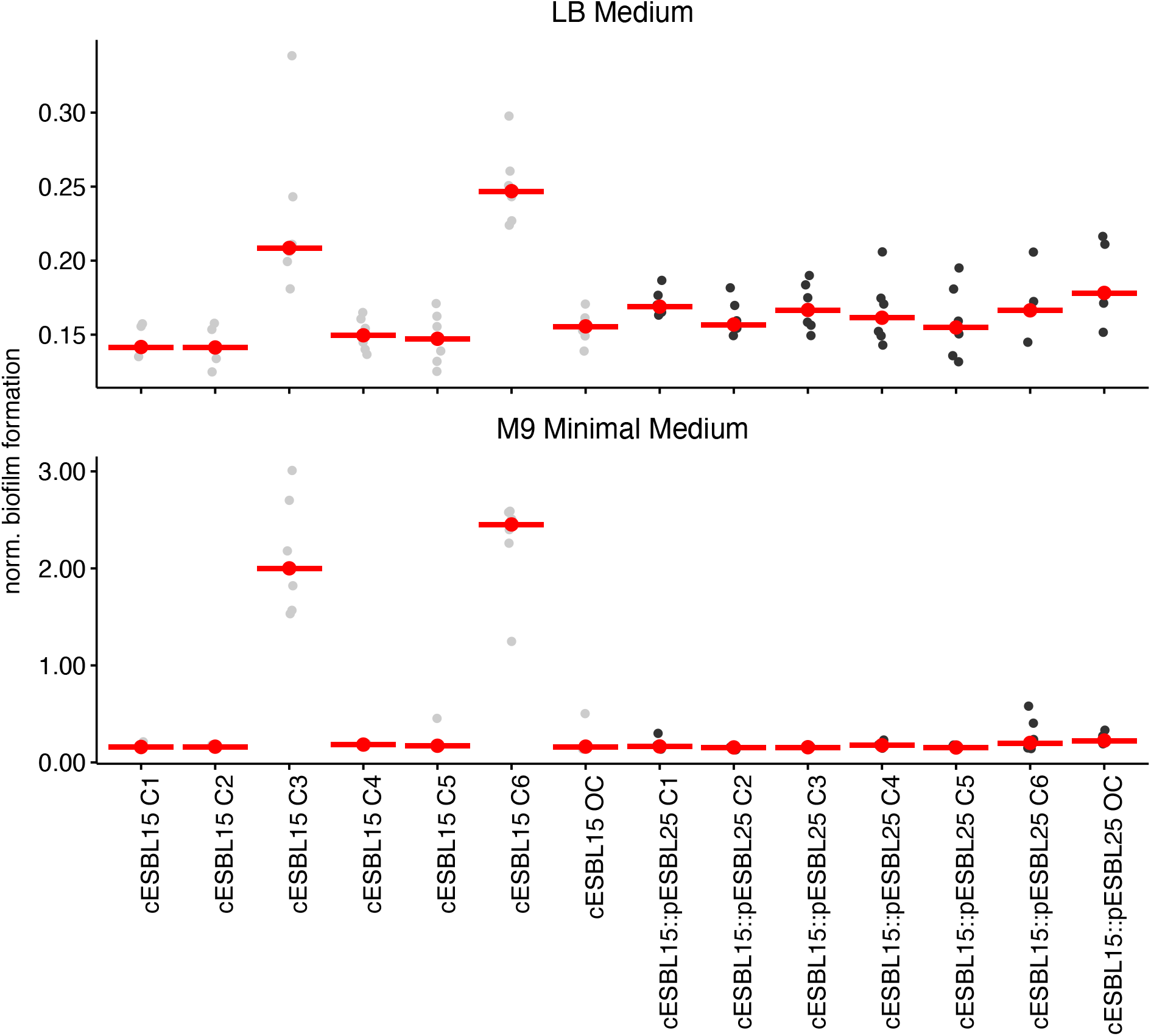
Normalised biofilm formation in independently generated clones (C1 – C6) of plasmid-free cESBL15 and cESBL15 with plasmid pESBL25. The six transconjugant clones were made as described in “Methods - Construction of novel plasmid-host combinations by conjugation”. The six independent control clones of cESBL15 were generated by exposure to the same treatment and culture conditions as the transconjugant clones were (i.e., cultured as if for a conjugation assay), but without adding the plasmid donor. We also included the original clones used in the previous biofilm experiment shown in the main text, labelled as cESBL15 OC and cESBL15::pESBL25 OC.

**Supplementary Figure 3:**
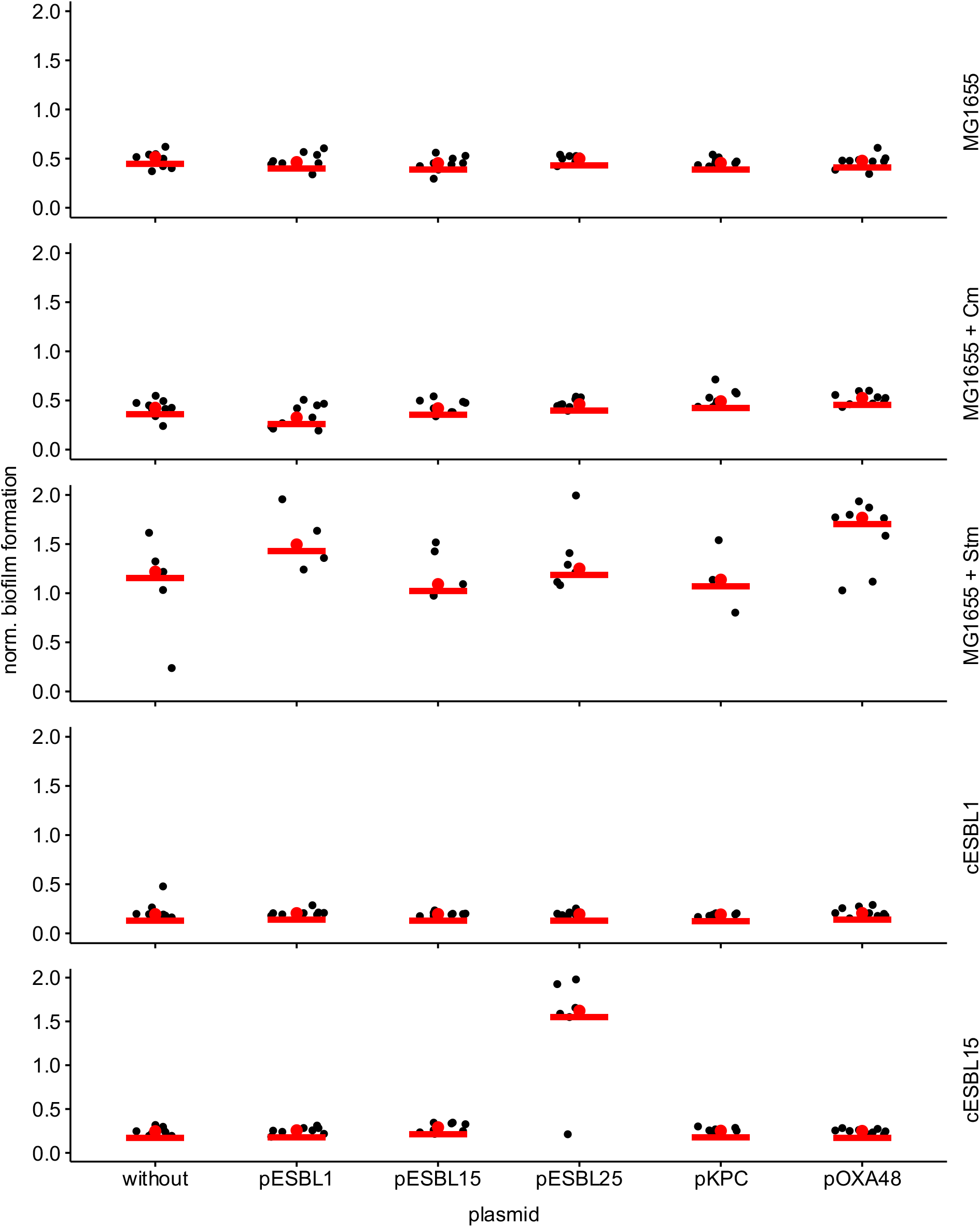
Normalised biofilm formation in M9 Minimal medium for 25 plasmid-bacterium combinations including plasmid-free reference strains. Each panel shows the biofilm formation in one host strain (labelled at right) with each of the five plasmids (x-axis). Data points show normalised biofilm formation in each replicate (n = 9).

**Supplementary Figure 4:**
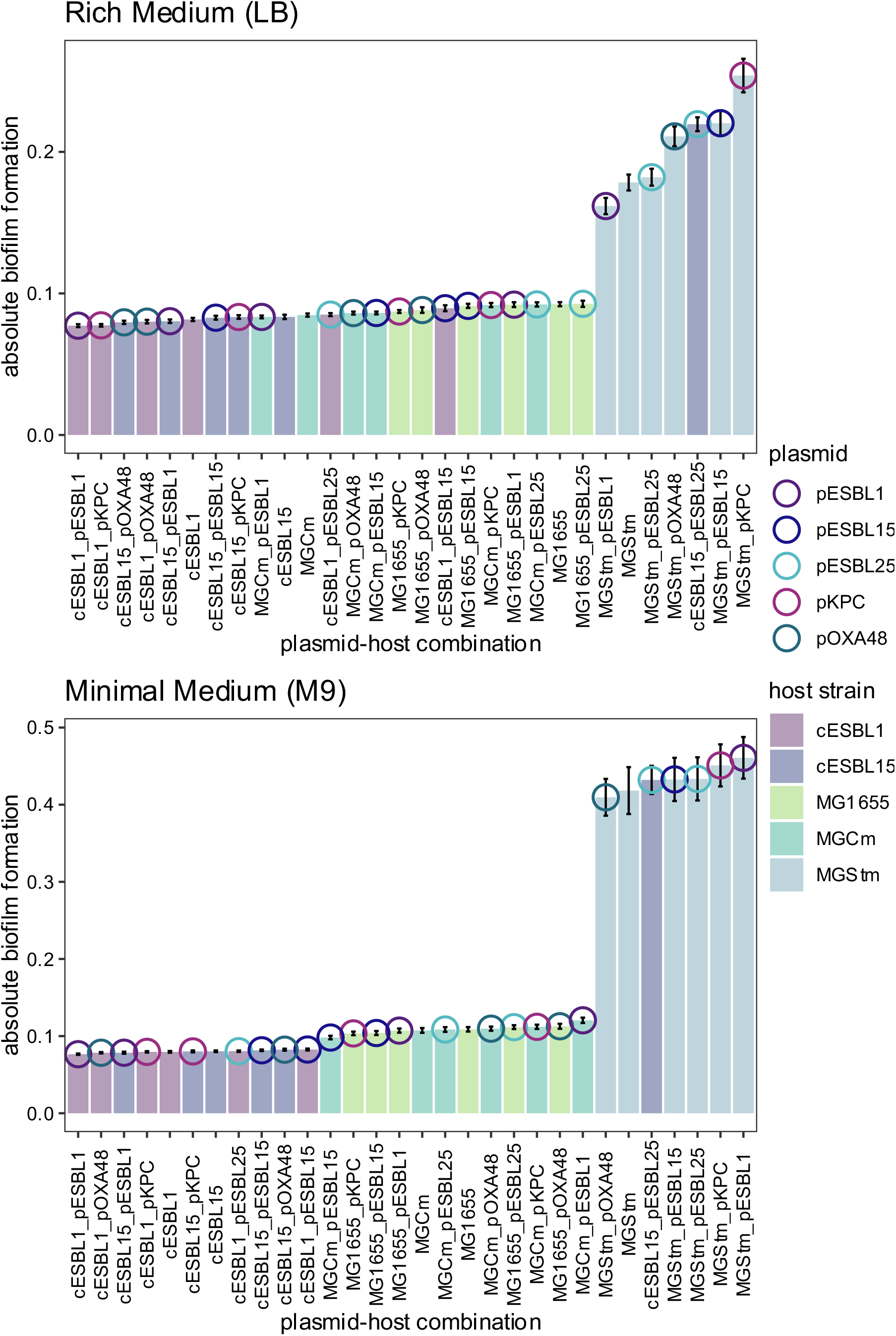
Absolute biofilm formation in LB medium (A) and M9 Minimal Medium (B). Absolute biofilm formation was quantified by measuring optical density after staining the adherent cells with crystal violet and dissolving in 95% EtOH (see Methods). Each bar represents the mean absolute biofilm formation for each strain (labelled on the x-axis by strain and plasmid in each combination; n=9 replicates per strain); error bars indicate propagated standard error.

**Supplementary Figure 5:**
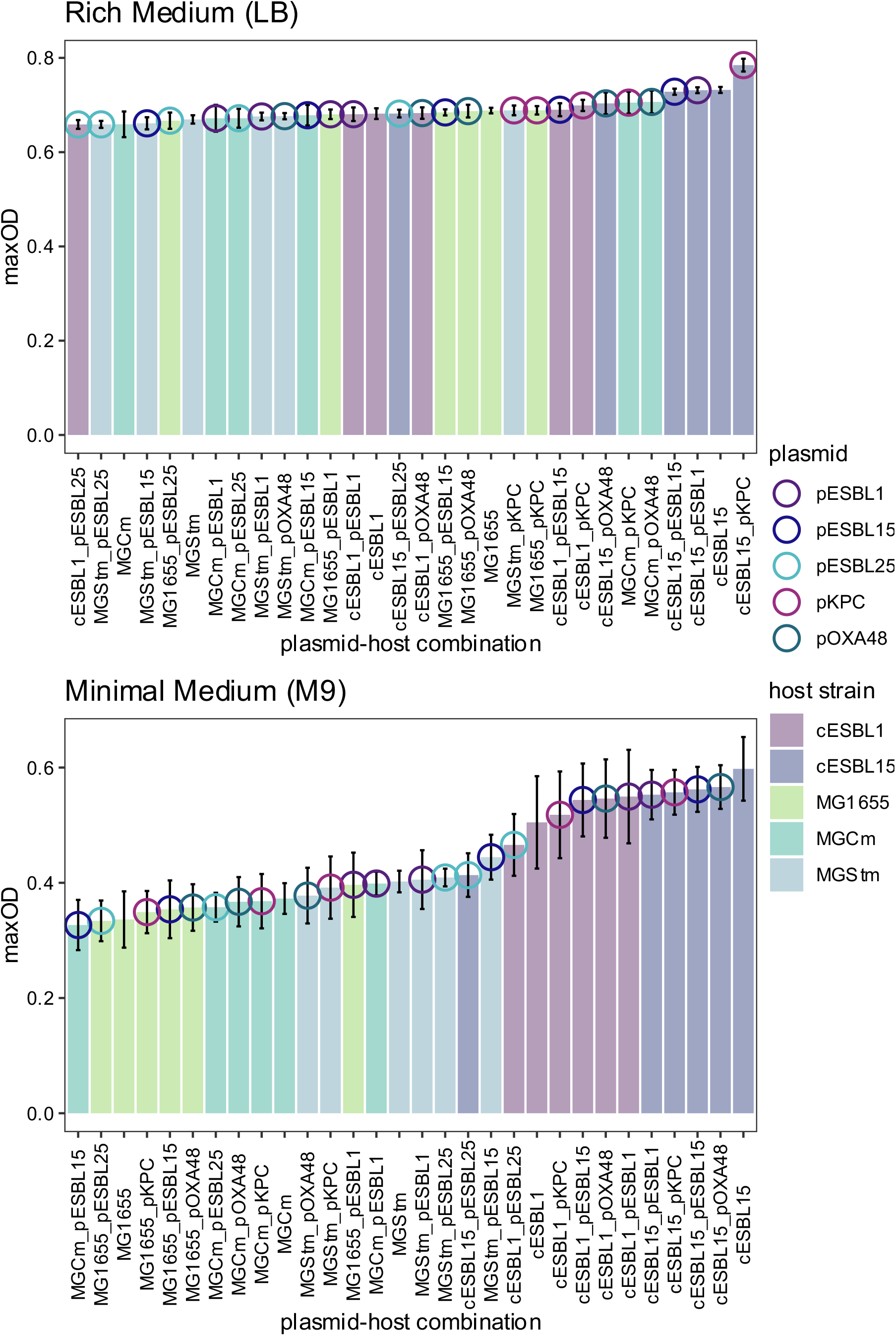
Maximum optical density (maxOD) in LB medium (A) and M9 Minimal Medium (B) for each plasmid-host combination. Each bar represents the maxOD for each strain (labelled on the x-axis by strain and plasmid in each combination; n=3 replicates per strain); error bars indicate propagated standard error. Two-way ANOVA effect of host strain: F = 13.34, p < 0.001, df = 4 (LB), F = 64.20, p < 0.001, df = 4 (M9); plasmid: F = 7.98, p < 0.001, df = 4 (LB), F = 5.23, p < 0.01, df = 4 (M9); and host:plasmid interaction: F = 1.42, p > 0.05 (n.s.), df = 16 (LB), F = 1.95, p < 0.05, df = 16 (M9).

**Supplementary Figure 6:**
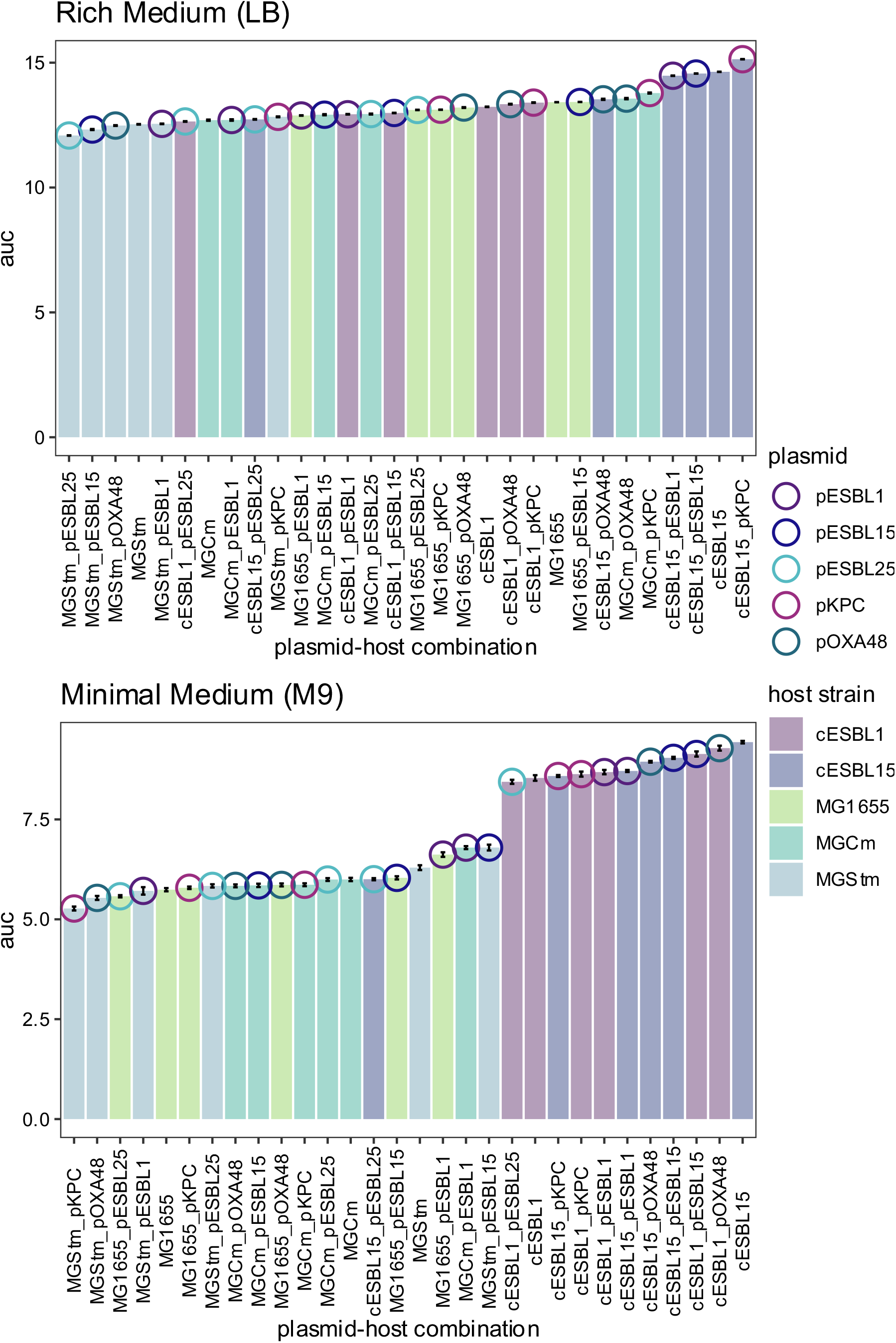
Area under the curve (AUC) in LB medium (A) and M9 Minimal Medium (B). AUC was calculated using the flux package in R. Each bar represents the auc for each strain (labelled on the x-axis by strain and plasmid in each combination; n=3 replicates per strain), error bars indicate propagated standard error. Two-way ANOVA effect of host strain: F = 21.62, p < 0.001, df = 4 (LB), F = 74.65, p < 0.001, df = 4 (M9); plasmid: F = 7.34, p < 0.001, df = 4 (LB), F = 6.02, p < 0.001, df = 4 (M9); and host:plasmid interaction: F = 2.39, p < 0.01, df = 16 (LB), F = 2.86, p < 0.01, df = 16 (M9).

**Supplementary Figure S2:**
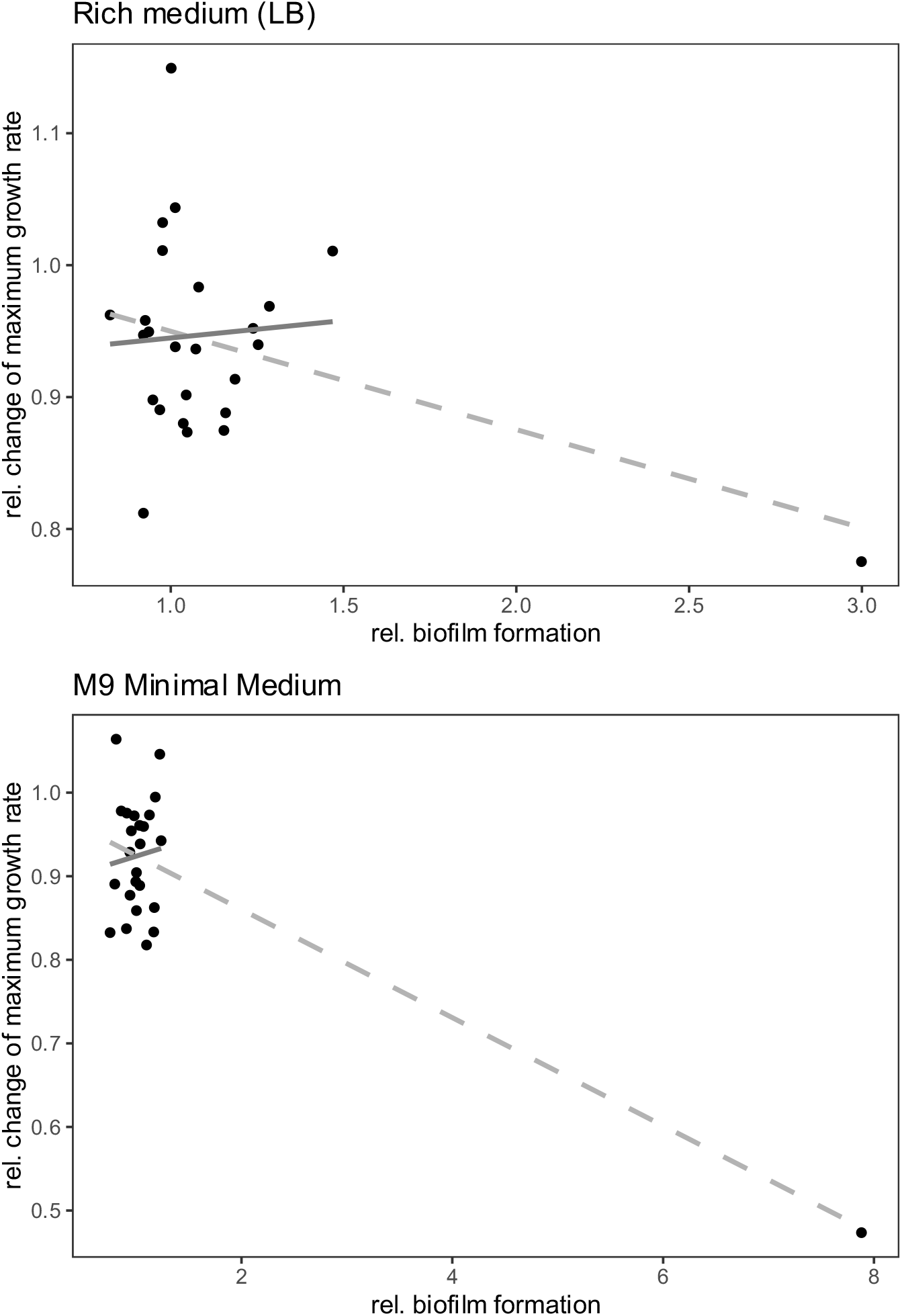
Correlation between plasmid biofilm effects and changes in maximum growth rate in LB and M9 Minimal medium. Linear regression models are shown in grey (dashed and light grey = whole data set, dark grey = subset excluding cESBL15::pESBL25), each data point represents one plasmid-host combination (n = 25). R2 = 0.12 (LB, whole data set) and 0.63 (M9, whole data set), R2= -0.04 (LB, subset) and -0.04 (M9, subset). Linear model coefficients for the whole data set in LB: intercept = 3.16 (p < 0.01), slope = -2.15 (p < 0.05) and in M9: intercept = 10.27 (p < 0.001), slope = -9.91 (p < 0.001). Linear model coefficients in the subset excluding cESBL15::pESBL25 in LB: intercept = 0.95 (p < 0.05), slope = 0.11 (n. s.) and in M9: intercept = 0.87 (p < 0.05), slope = 0.16 (n. s.).

### Supplementary Tables

**Supplementary Table 1:**
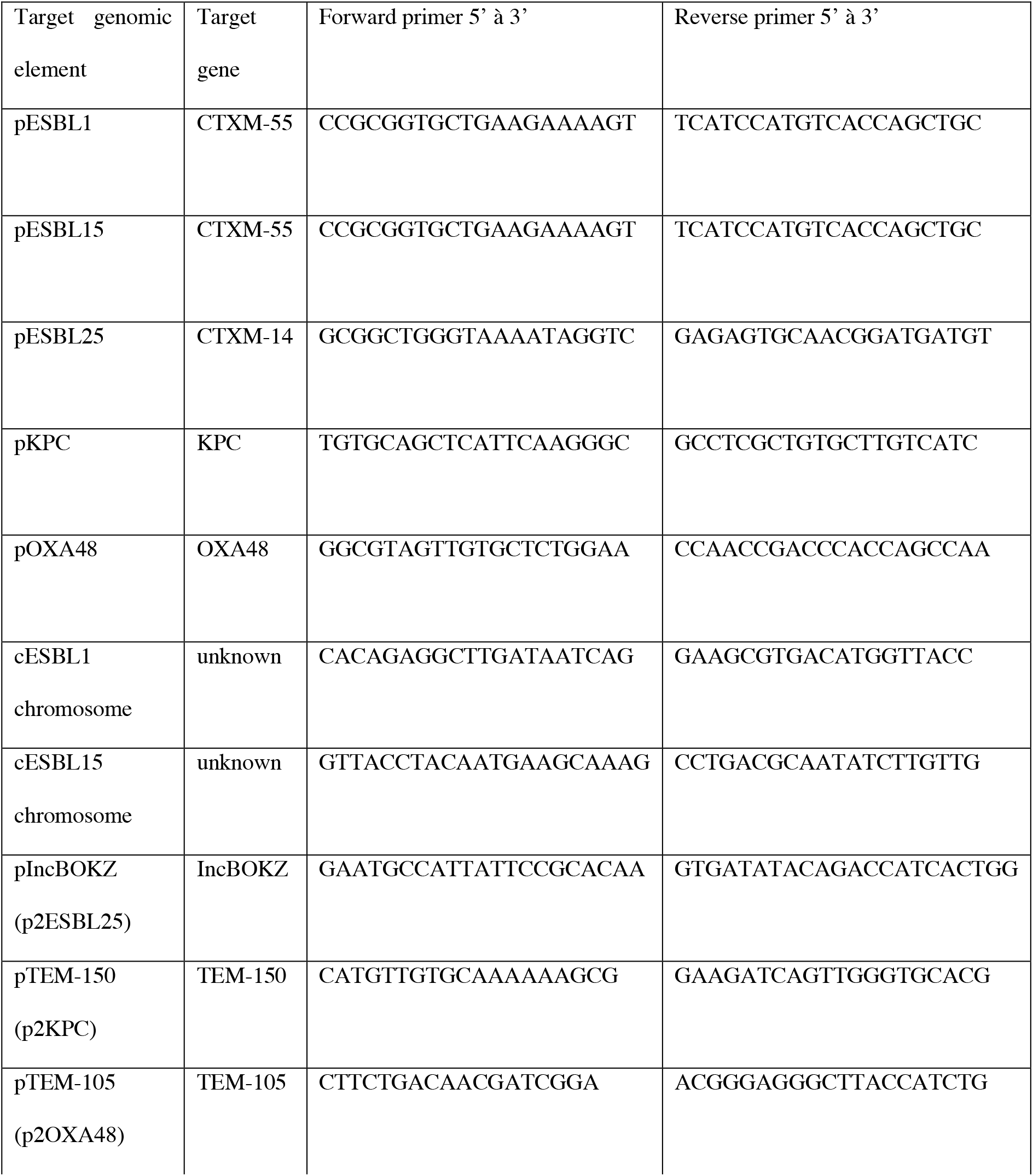
Primers used for colony PCR reactions.

**Supplementary Table 2:** Putative entry exclusion system genes associated with each plasmid family studied were identified aligning the plasmid sequences against the reference sequences.

| Plasmid | Incompatibility group | Entry exclusion system genes | Nucleotide position in the plasmid | Reference protein ID |
| --- | --- | --- | --- | --- |
| pESBL1 | IncI | *excA*, *traS*, *traG* | *excA*: 64117-64767;<br>*traG*: 86220-86804;<br>*traS*: 73420-73608 | *excA*: WP_001178506.1;<br>*traG*: WP_000977522.1;<br>*traS*: WP_001277255.1 |
| pESBL15 | IncI | *excA*, *traS*, *traG* | *excA*: 45400-46050;<br>*traG*: 67498-68082;<br>*traS*: 54697-54885 | *excA*: WP_001178506.1;<br>*traG*: WP_000977522.1;<br>*traS*: WP_001277255.1 |
| pESBL25 | IncFIB, IncFIA | *traS*, *traG* | *traS*: 109875-110384;<br>*traG*: 107101-109878 | *traS*: WP_000628111.1;<br>*traG*: WP_152930645.1 |
| pKPC | IncX3, IncU | *traG* | *traG*: 49001-50836 | *traG*: VCY81245.1 |
| pOXA48 | IncL | *traY*, *excA* | *traY*: 60121-62316;<br>*excA*: 62318-62971 | *traY*: AEV46259.1;<br>*excA*: AEV46260.1 |

## Notes

### Competing Interest Statement

The authors have declared no competing interest.

